# Beyond biomass: how interactions shape species’ role for ecosystem functioning

**DOI:** 10.1101/2025.06.11.659013

**Authors:** Alice N. Ardichvili, Michel Loreau, Thomas Onimus, Nuria Galiana, Ismaël Lajaaiti, Sonia Kéfi, Jean-François Arnoldi

## Abstract

Assessing the functional role of species in a changing world is critical for effectively preserving ecosystems. Intuitively, this role should relate to how the loss of a species ultimately impacts a function. We introduce the notion of a species dynamic contribution, which takes into account biotic interactions as they develop in the community. Contrary to the static contribution —a measure of what a species directly does— the dynamic contribution captures what the presence of the species causes. Using both model simulations and empirical data from a ciliate microcosm experiment, we demonstrate that dynamic and static contributions of species are generically unrelated. Our novel characterization of species’ functional contributions reveals that, due to biotic interactions, rare species that do not possess unique functional traits are just as likely as any other species to play an important role for ecosystem functioning.

## Introduction

Understanding how ecosystems respond to global change is a central goal of functional ecology and conservation biology (Lavorel et al. 2002, Hong et al. 2022). In the context of the ongoing biodiversity crisis (IPBES, 2019), assessing how changes in community composition influence ecosystem functioning is crucial and requires a deeper understanding of the roles species play in maintaining ecosystem functions.

Two important fields of research, functional ecology and Biodiversity and Ecosystem Functioning (BEF) have tackled species’ roles with different approaches and conclusions. Functional ecology posits that knowledge on traits at the species level (eg. leaf area index, photosynthetic rate) captures the mechanisms structuring biodiversity and ecosystem functioning (Enquist et al., 2015; Lavorel and Garnier, 2002). The mass-ratio hypothesis introduced by Grime (1998) further suggests that the immediate effect of a species on an ecosystem function is proportional to its biomass, in which case the functioning of ecosystems is expected to be largely driven by the traits of dominant species. Accordingly, community-weighted trait means obtained from a snapshot of a community have generally been used to predict the impact of global change on ecosystem functioning (Garnier et al., 2004).

The BEF research field has consistently shown that species richness is a strong determinant of ecosystem functioning regardless of species identity, because species with different traits tend to occupy different niches and therefore to be functionally complementary to each other (Loreau et al. 2001; Loreau & Hector 2001; Hooper et al. 2005; Cardinale et al. 2007; Loreau et al. 2022). Results from BEF research imply that the contribution of species to ecosystem functioning largely depends on the ecological context in which species live, in particular on how they interact and partition resources. As a result, determining which species have a significant impact on ecosystem functioning is challenging (but see Jaillard et al., 2018). While the mass-ratio hypothesis and BEF approaches are often opposed (eg. Wasof et al. 2018; Smith et al. 2020; Brun et al. 2022), they are not necessarily contradictory, but a unifying perspective that assesses the importance of species in their ecological context is still lacking.

The contrast between the mass-ratio hypothesis and BEF is blatant when it comes to assessing the role of rare species. Under the mass-ratio hypothesis, rare species are assumed to generally play a minor role in ecosystem functioning. From the BEF perspective, species richness itself matters even for broad functions (eg. biomass production). Rare species in BEF assemblages should therefore be of some importance to ecosystem functioning. Supporting this perspective, *in situ* and *in silico* removal experiments have shown that removing rare species can disproportionally affect ecosystem functioning (Heilpern et al. 2018; Delalandre et al., 2022; Leitão et al. 2016; Peltzer et al., 2009).

A key yet implicit difference between the mass-ratio hypothesis and BEF approaches lies in the role given to biotic interactions. The mass-ratio hypothesis focuses on the biomass distribution in a community at a given point in time, without considering biotic interactions and the resulting ecological dynamics. A dominant species contributes the most to the total biomass of a community (or any other broad function), but its presence could -via competitive interactions-suppress the growth of otherwise more productive species. A species that is rare today may have played a key role in the past, facilitating the establishment of a productive species (Law and Morton, 1996; Koffel et al., 2018). In the BEF literature, a strong, positive selection effect (*sensu* Hector and Loreau, 2001) implies that dominant species perform most of the ecosystem function, a view that coincides with the mass-ratio hypothesis. However, BEF experiments have shown that the complementarity effect generally predominates, while the selection effect is often variable, weak on average, and decreases over time (Loreau & Hector 2001; Cardinale et al., 2007; Fargione et al., 2007). The contributions of species to ecosystem functioning, if seen in a general sense that accounts for the ecological dynamics, cannot be understood without explicitly considering species interactions.

Interactions between species result in indirect effects, feedback loops, and cascading events (Pichon et al., 2024; Harvey et al. 2017). Although increases in interaction complexity leads to higher context-dependency (Zelnik et al., 2024), this does not necessarily imply that all ecosystem responses become unpredictable (Lajaaiti et al., 2025; Liautaud et al., 2019). Recent experimental findings by Sanchez et al. (2023) demonstrate that the functional impact of a species invasion can often be predicted from basic knowledge of the biotic context in which the species is added. How to link the existence of such predictable relationships to interaction patterns remains an open question, but one that must be addressed if we want to understand the functional role of species within a community.

The purpose of this article is to reveal how biotic interactions shape the functional role of species in a community. In functional ecology, the role of species is captured by a partitioning of functions into species-level contributions that takes into account their functional (effect) traits weighed by their biomass (Garnier et al., 2004). We call this the **static contribution** of species. Species interact however, such that they will affect ecosystem functions beyond their static contributions. Our first objective is to define a partitioning of ecosystem functions into **dynamic contributions** of species. This partitioning should take into account species direct and indirect interactions as they develop in the community, and as such should leave a recognizable signature in the response of ecosystem functions to certain biotic or abiotic perturbations. Our second objective is to compare static and dynamic contributions of species, using both simulated and empirical communities from a ciliate microcosm experiment (Pennekamp et al., 2018a). Our third objective is to understand how a species’ interaction pattern can make its dynamic contribution predictably context-dependent, or on the contrary, highly sensitive to details of the community structure in which it grows.

## Methods

### Defining a dynamic partitioning of an ecosystem function

To meet our first objective, we start from the mass-ratio hypothesis in which each species contributes to a function Φ via its biomass *N*_*i*_ [g] and its functional contribution per unit of biomass *t*_*Ei*_ (as in Lavorel and Garnier, 2002; Violle et al., 2007; Heilpern et al., 2018). For the sake of conciseness, we call *t*_*Ei*_ ‘effect trait’ but we may keep in mind that this quantity, which converts biomass of species into the unit of the function Φ, reflects the species-level expression of potentially several phenotypic traits (Diaz, 2025). This idea translates as the following linear decomposition:

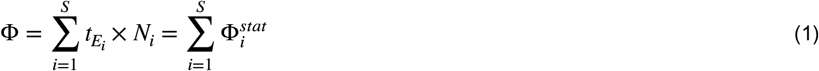

In this partitioning of the ecosystem function, each species has a **static contribution** 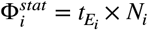 which is observable at any moment in time, from a snapshot of the community. This decomposition applies to different functions (eg. total biomass, nitrogen litter content, polymer degradation) as the units of effect traits change (Fig. 1b). When the function is total biomass, all species have the same contribution per unit of biomass, and the unit of the function is the same as species biomass; all effect traits are 1 (dimensionless). If the function is phosphorous litter content [mg g^−1^], the effect traits capture the species-specific amount of litter produced per unit of biomass and the phosphorous concentration of the produced litter, i.e. in units [mg g^−2^]. In what follows, we assume that the effect trait only depends on species identity. Although effect traits can be more or less density dependent (metabolism, Fant and Ghedini, 2024; enzyme production, Sanchez-Gorotsiaga et al., 2019), assuming constant effect traits is a useful starting point for our theory, and one that reflects the assumptions underlying the use of community weighted means in functional ecology.

**Figure 1.**
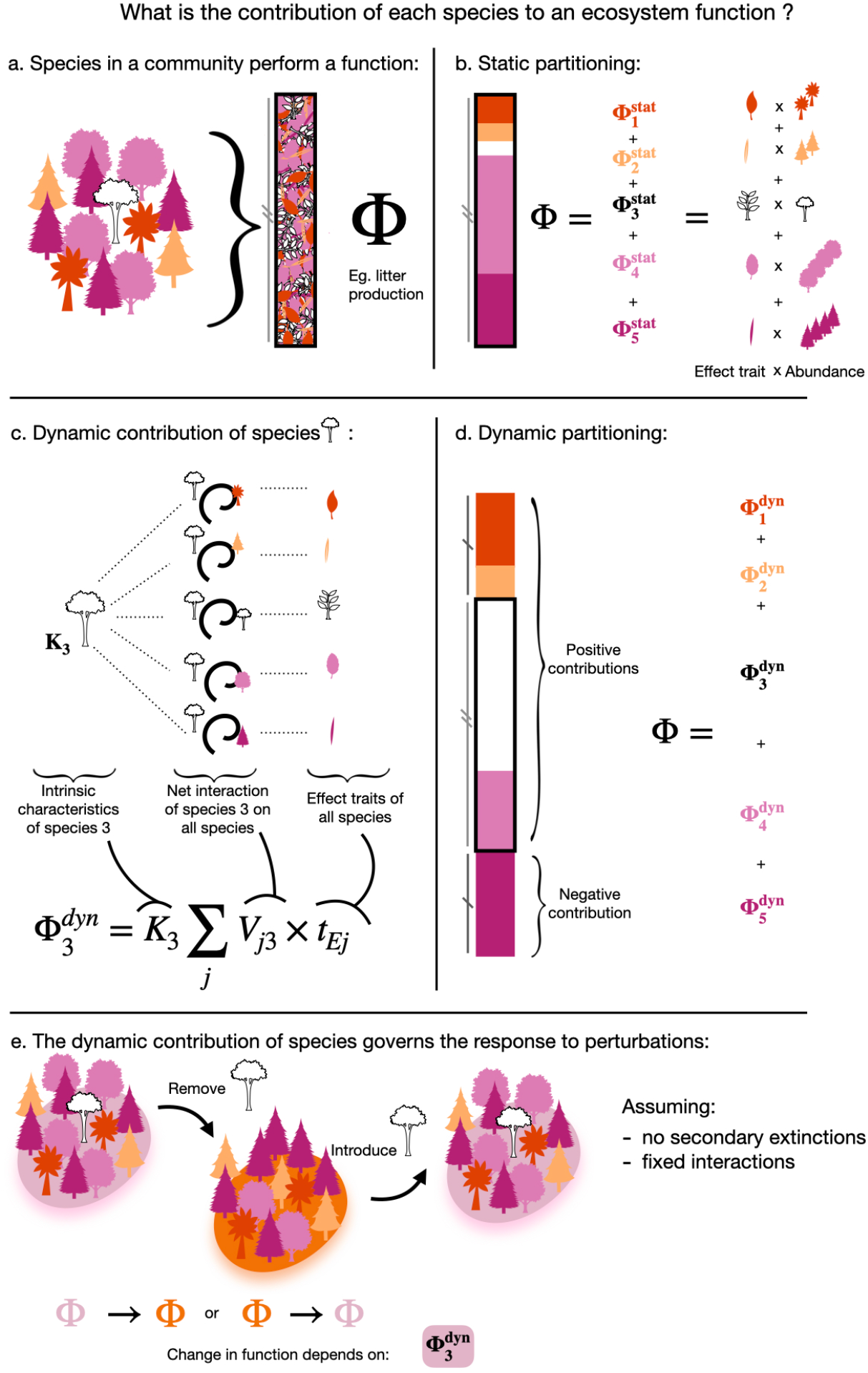
An ecosystem function can be partitioned into static (b) and dynamic (d) contributions of species. While the static contribution depends only on the traits and abundance of a given species, the dynamic contribution of one species links the effect trait of every other species to the intrinsic characteristic of the focal species via net interactions (c). The dynamic contribution captures the response of ecosystem functioning to perturbations such as extinctions or introduction of species (e).

The partitioning in Eq. 1 only *implicitly* considers the role of species interactions, which come into play as ecological dynamics unfold over time. Interactions between species influence their biomass, and thus their static contribution to functions. Our goal is to define a notion of functional contribution of species that *explicitly* accounts for ecological dynamics. A key trick is to introduce the matrix *V*_*ji*_ = ∂*N*_*j*_ /∂*K*_*i*_ which represents the net interaction (sum of direct and indirect interactions) from species *i* to species *j* (Zelnik et al. 2024), and define 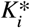 such that 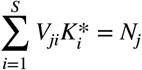. Note that for Generalised Lotka-Volterra (GLV) models, at equilibrium, we have 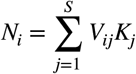 so that 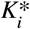 is exactly the carrying capacity of species *i, K*_*i*_. Here we use a notation referring to carrying capacities 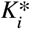 but it could also refer to growth rates or any parameter reflecting a species performance when alone in the environment. Deviations between 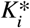 and *K*_*i*_ occur for community models departing from GLV dynamics or from equilibrium conditions (Appendix C). The partition of the ecosystem function (Eq. 1) can therefore be written as:

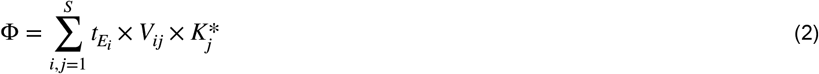

Equation 2 shows, with the summation over the two indices *i* and *j*, that the function Φ can be partitioned in two different ways. Summation over *i* represent the static partitioning; the summation over *j* reveals a dual partition that we call the **dynamic contributions** of species:

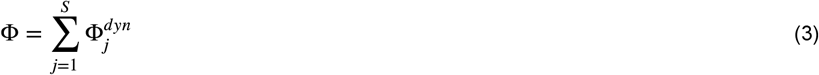

where

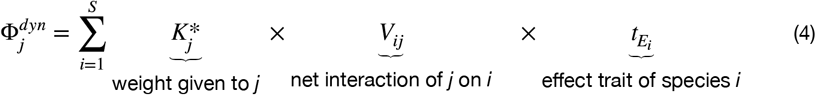

Equation 4 shows that the dynamic contribution of a species *j*, 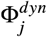, does not only depend on that species effect trait (Fig. 1c). For example, 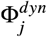 can be large if species *j* has a positive net interaction on other species with large effect traits. On the contrary, the dynamic contribution of species *i* decreases if it has a negative net interaction on species with strong effect traits. The static and dynamic contributions of species possibly have different values, and even different signs since the dynamic contribution of a species can be negative.

Because 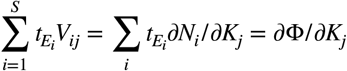, by rearranging the terms of Eq. (4), we show that this dynamic contribution reflects the sensitivity of a function to changes in intrinsic demographic parameters of species:

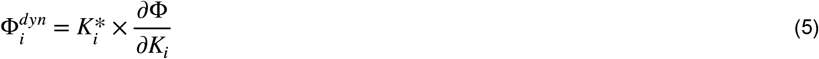

Equation 5 means that the dynamic contribution of species encodes the response of the function to a linear press perturbation of the species’ intrinsic demographic parameter 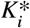. The perturbation could be caused by biotic (eg. species invasion or extinction) or abiotic (eg. pollution or habitat destruction) disturbances. Here we assume that interactions remain constant (as would arguably be the case for eg. habitat destruction or species introduction), but even if they do change, we show in appendix E that the above formalism still holds, because a change of interactions can be still be seen as an effective change in the environment (and thus formalized by a change in parameter 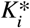). The important take away here is that the dynamical contribution of species is inherently related to patterns of ecosystem response to perturbations. This means that we can use response to perturbations to infer species dynamic contributions. This is precisely what we do in the next section, using data from a BEF experiment..

### Contrasting the dynamic and static contributions in empirical and simulated communities

Our next objective is to show that the static and the dynamic contribution are generically uncorrelated. To infer the dynamic contribution from empirical data, a next step is necessary. We consider a targeted press perturbation that leads to the extinction of the focal species *i*: *K*_*i*_ → *K*_*i*_ + *δK*_*i*_ such that *V*_*ii*_*δK*_*i*_ = − *N*_*i*_ (Appendix A, Eq. A3). Let ΔΦ_*i*_ = Φ − Φ^\*i*^ be the change in ecosystem functioning due to the removal of the species *i* from the community. We define this difference so that a positive value implies that the species has a net positive impact on the function in question, and thus that its removal is detrimental. We apply our dynamical partitioning (Eq. 5) to predict the effect of the perturbation *δK*_*i*_ = − *N*_*i*_ /*V*_*ii*_ that pushes the species to extinction. Assuming that the extinction of species *i* does not cause secondary extinctions (assumption relaxed in Appendix D), the effect of removing a species is directly linked to its dynamic contribution:

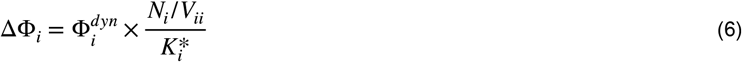

The term *V*_*ii*_ encodes the species sensitivity (Arnoldi at al 2022), and is uniform across species for disordered networks or when interactions are weak (Bunin 2017). Conversely, this means that the dynamical contribution of a species can be estimated from the effect of its removal in a given community:

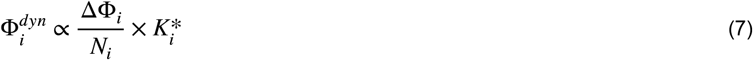

where the proportionality constant is the mean sensitivity. Since the sum of 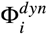 is known (=Φ), so that dynamical contributions are relative notions to begin with, we do not need to know what this mean sensitivity precisely is.

With this approximation of the dynamic contribution of a species at hand, we analyzed data from an experimental ciliate community experiment (Pennekamp et al., 2018b). Six species (*Tetrahymena, Colpidium, Dexiostoma, Loxocephalus, Paramecium, Spirostomum)* were grown at 53 different diversity levels during 57 days, with complete representation of the levels of richness 1, 2, 5 and 6. We focus on total biomass for which the effect trait of every species is 1. We show results from communities grown at 19°C.

#### Estimating biomass and interactions

After visual inspection of the time series of species abundances, we used the last 6 data points of the experiment, at which the biomasses approximately stabilized, and retrieved the mean biomass of each species in each community. We equated 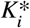 to the abundance that species *i r*eaches in isolation, and obtained the pairwise interactions from duo cultures and monocultures, such that 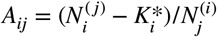. For all six species, we estimated their mean interaction effect 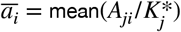.

#### Obtaining the change in ecosystem function

We measured the change in total biomass due to the removal of a species by comparing pairs of communities which only differed by one species in their composition. For each species, we obtained 21 comparisons, yielding a total of 126 comparisons for the 6 species.

*Simulations*: We ran simulations of a GLV model with *S=9* species, where parameters were randomly drawn to showcase the genericity of our results. We integrated the differential equations with the DifferentialEquations package (v1.9.0, Rackauckas and Nie, 2017) in Julia (v1.10.4, Bezanson et al. 2017), until an equilibrium was reached (biomasses did not vary by more than 10^−6^ between two integration time steps). The carrying capacities and effect traits were drawn from a uniform distribution between 0.5 and 1 and 0 and 1 respectively. The mean interaction strength 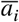 was drawn between −0.9 and 0.5, and the deviation from mean interaction Δ*a*_*ij*_ drawn between *u*_*a*_ and *u*_*a*_, so that parameter *u*_*a*_ governs the heterogeneity of interactions (*u*_*a*_ close to 0 implies low variability in interactions while a large *u*_*a*_ implies variable interactions).

### Understanding how the pattern of a species’ interactions with others relates to its dynamic contribution

Our final objective is to demonstrate that the context-dependence of a species’ dynamic contribution is shaped by patterns of species interactions. Two key aspects of these interaction patterns are the average (mean) and variability (variance) of a species’ effect on others. We will show that simple relationships can emerge between a species’ dynamic contribution and the overall functioning of the community in its absence — a baseline we refer to as the **background level of functioning** (Φ^\*i*^), following Sanchez et al. (2023).

The dynamic contribution of a species 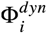 reads (see Appendix A for derivation):

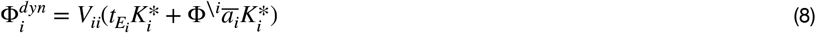

In this expression, 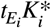 is the level of ecosystem functioning in monoculture. The second term in brackets encodes context-dependency via the background level of functioning Φ^\*i*^. The term 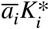 can be interpreted as the weighted mean ofthe interactions incoming from species *i*. This weighted mean has to be understood with care since the weights are the relative dynamical contributions of affected species, and those contributions could be negative. If the interactions *a*_*ki*_ are homogeneous (i.e. not variable), the dynamic contribution of a species depends on its average effect on other species. A species that tends to facilitate all other species therefore has a positive dynamic contribution, and its contribution increases as the overall ecosystem functioning increases. A species that competes homogeneously with all other species has a declining dynamic contribution as the overall level of ecosystem functioning increases, such that its dynamic contribution may end up negative. If the interactions are heterogeneous, an apparently facilitative species (i.e. one that facilitates most species present in the community) can have a negative effect on functioning if it facilitates a strongly antagonistic species (i.e. a species whose dynamical contribution is negative).

From Eq. 8, we therefore expect that when a species has an homogeneous effect (with a low variance) on others, its dynamic contribution should be linearly positively or negatively linked to the level of background functioning depending on the mean interaction effect. On the contrary, we expect that when a species has an heterogeneous effect (with a high variance) on others, its dynamic contribution is independent of background functioning. We test whether this relationship exists in empirical and simulated communities.

## Results

In the section **Defining a dynamic partitioning of an ecosystem function**, we defined a partitioning that reflects species direct and indirect the indirect contributions to a given function. In a community, the sum of the dynamic contributions of all its constituent species add up to the ecosystem function (Objective 1).

In figure 2, we contrast the static and dynamic contributions of all species across biotic contexts (Objective 2), showing that they are not correlated in both simulations (Fig. 2a) and empirical data (Fig. 2b). On the right, dots correspond to species that are dominant (they have a high static contribution), but their vertical position shows that these species can both have a null dynamic contribution (all dots close to the black line, *Paramecium, Spirostomum* and *Colpidium* in 2b), negative contributions (eg. *Colpidium* and *Paramecium* in two communities in b) or a strong positive contribution (eg. yellow and orange species in 2a). On the contrary, on the left, dots correspond to species that are rare or have a low effect trait, but their vertical position indicates that their dynamic contributions are strongly positive (eg. pink species in 2a, *Loxocephalus* in b) or negative (black species in a, *Dexiostoma* in 2b).

**Figure 2.**
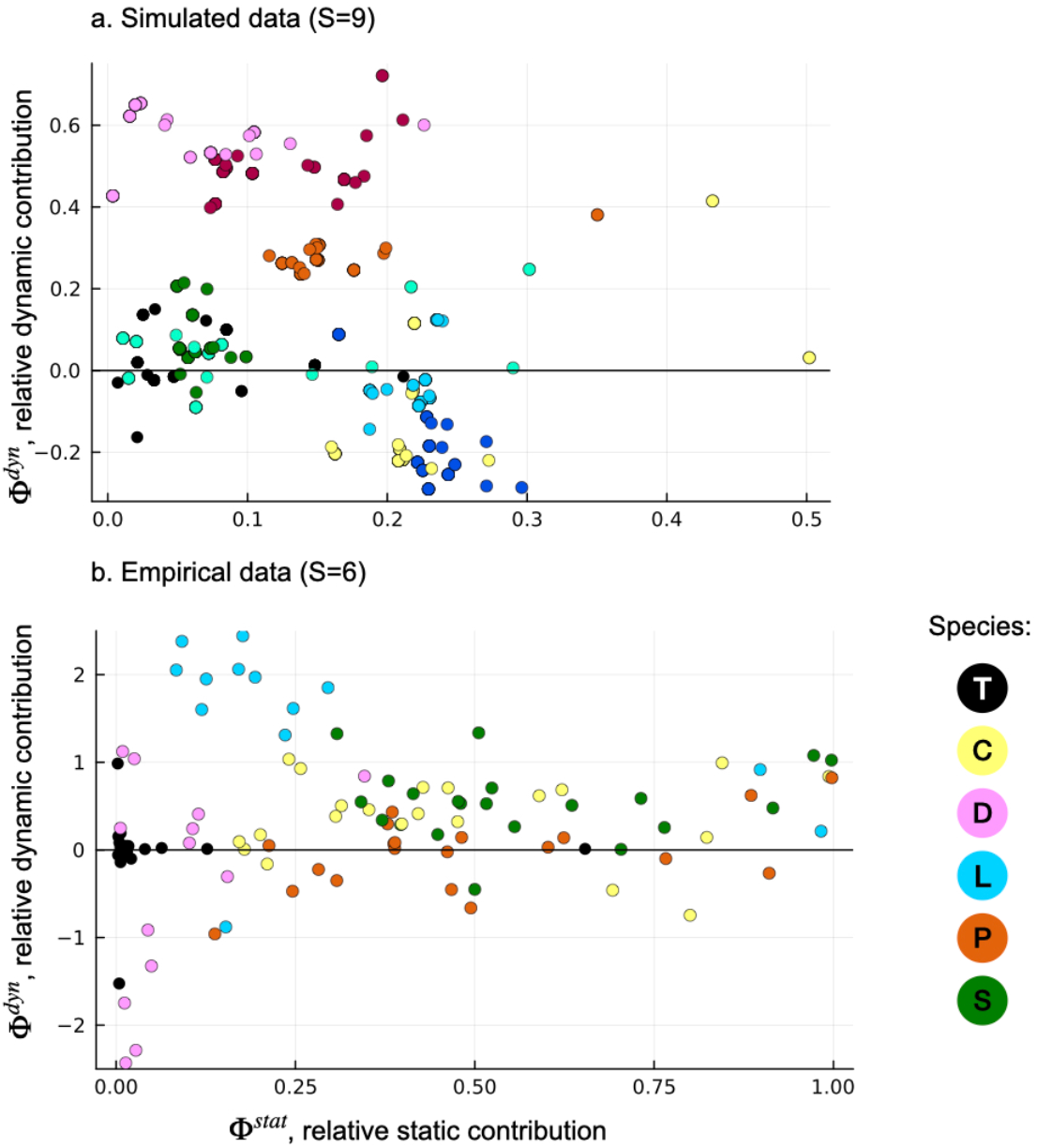
Dynamic contributions are not necessarily correlated to static contributions. Each dot corresponds to a species in a community, dots are colored by species. There are species that tend to have a low static contribution but a high dynamic contribution (eg. pink species in a, *Loxocephalus* (blue) in b), species with a high static contribution but a low dynamic contribution (eg. yellow species in a, *Paramecium, Spirostomum and Colpidium* (orange, green and yellow) in b), and species whose dynamic contribution vary in sign (eg. black and aquamarine species in a, *Dexiostoma* (pink) in b)

In figure 3 and 4, we illustrate how species’ interaction patterns (mean and variance of its effect on others) are reflected in their dynamic contributions across biotic contexts (Objective 3). Specifically, we show that background functioning and the mean effect of a species on others explains its dynamic contribution, but only when the variance in the effect that that species has on others is low enough. Figure 3 illustrates the effect of the variance in simulations: when the variance of interactions is low (Fig. 3a), the dynamic contribution of species is explained by the level of background functioning and by the mean effect of that species (see how dots fall on the line). As the variance in interactions increases (Fig. 3b), the relationship between the background functioning and the dynamic contribution fades. In the empirical data (Fig. 4), the relationship is strong for species *Colpidium, Paramecium*, and *Spirostomum*, which have the lowest variance in their effects on others (*σ*_*a*_ are low in Fig. 4). By contrast, *Tetrahymena, Loxocephalus*, and *Dexiostoma*, whose interactions are much more variable (see how on a single column of Fig. 4g, positive and negative values exist; this column-wise variability is reflected by high *σ*_*a*_ in Fig. 4), showcase idiosyncratic dynamical contributions across biotic contexts, unrelated to background functioning. Our theory explains these inconsistent contributions as the result of complex indirect effects within the community where the species are introduced, resulting from high variability in species interactions.

**Figure 3.**
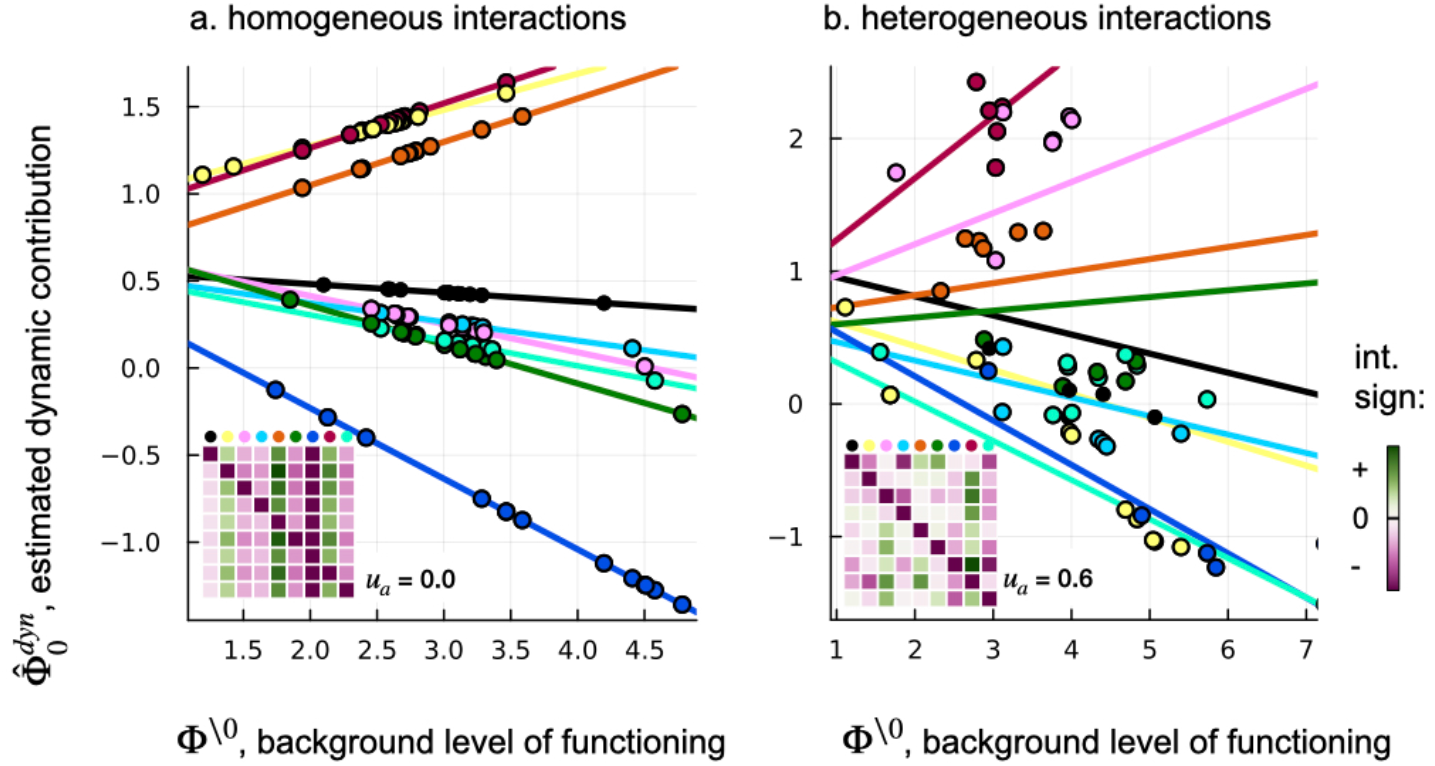
Estimated dynamic contributions of species against background functioning in two simulated communities. Every dot corresponds to a species in one community comprised of a subset of the 9 species. The solid line is the expected relationship between the dynamic contribution and expected functioning. The inset matrix represents pairwise interactions in the simulated community; column wise, you can see the effect that each species (small dot at the top of the column) has on others. When the effect of the species on others is homogeneous (*u*_*a*_ = 0, see how columns of the interaction matrix are of the same color), its dynamic contribution scales linearly with the level of functioning prior to the introduction Φ^\0^ (panel a). However, when the introduced species has an heterogeneous effect on others (*u*_*a*_ = 0.6), the relationship between the dynamic contribution and the background ecosystem functioning erodes (panel b).

**Figure 4.**
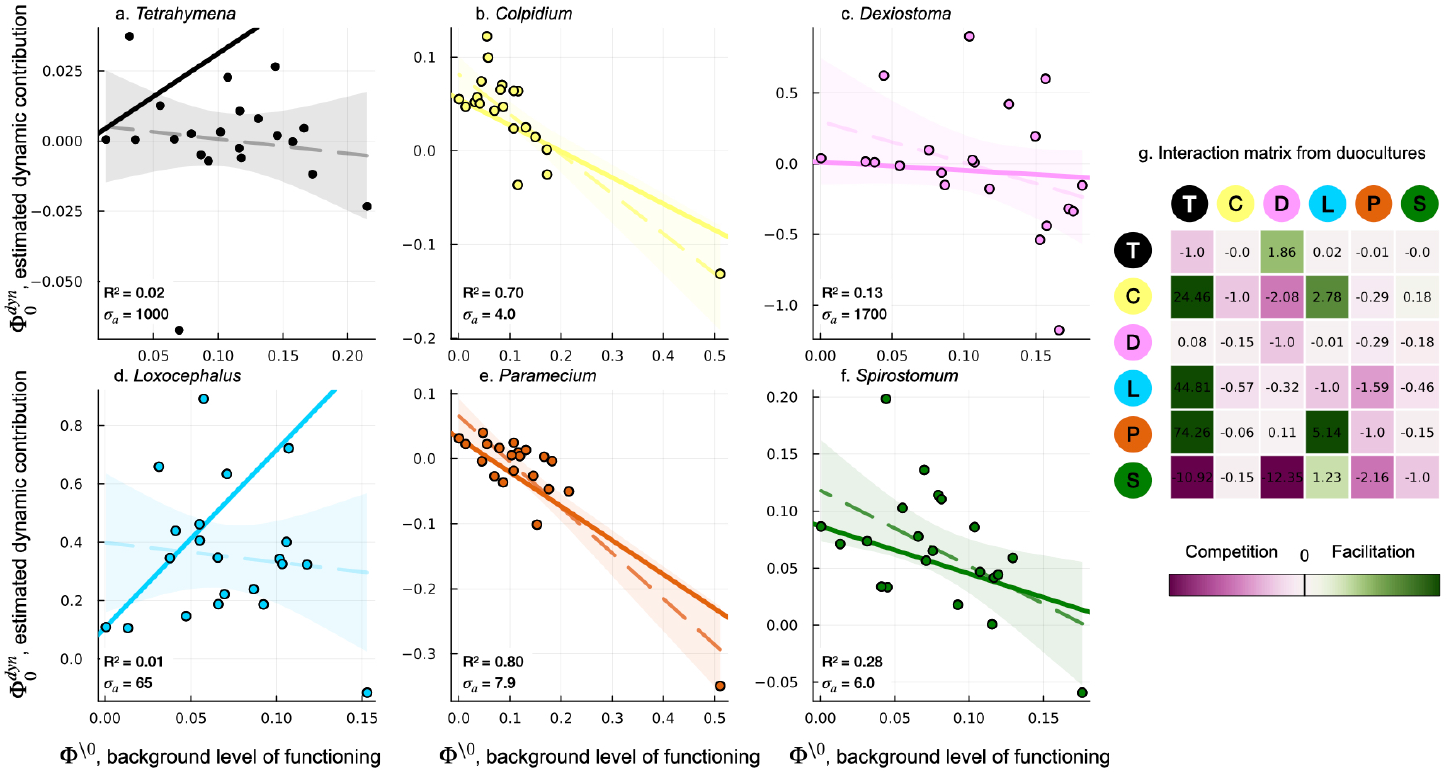
The dynamic contribution of some species is related to the background level of functioning, via the species’ mean effect on other species. a-f: The solid line is the theoretical expectation from Eq. 8, while the dashed line is the statistical relationship between background functioning and the dynamic contribution, and the associated R^2^ is on the bottom left of the panel. g: direct interactions (unscaled) between the 6 species in the communities. These direct interactions are scaled by the recipient species’ carrying capacities to obtain the theoretical expectation in panels a-f. For 3 species, the statistical relationship closely aligns with the theoretical relationship (panels b, e and f). These species happen to have the most homogeneous effects on others, as shown by the low standard deviation in felt interaction *σ* _*a*_ of the three species. For the other three species, the dynamical contribution seems unrelated to the background level of functioning.

## Discussion

We introduced a partitioning of any ecosystem function into species-level contributions in a way that fully takes into account the interactions between those species. This partitioning is reveals the role played by species in the ecosystem-level response to certain perturbations, whose impact on individual species and interactions are known (a species removal being arguably the simplest experimentally accessible example of such perturbations). Contrary to the static contribution derived from the mass-ratio hypothesis —a measure of what a species directly does— the dynamic contribution captures what the presence of the species *causes*. This contribution is context dependent by construct, but when a species has a relatively homogeneous effects on other, its dynamical contribution is nonetheless predictable, and follows a simple rule that connects the intrinsic characteristics of the focal species to the ecological context in which it grows. In any case, within a community and because of biotic interactions, the static and dynamic contributions of a species are generically uncorrelated. This explains how the mass-ratio hypothesis can be a valid static description of the role played by species in an ecosystem and yet be misleading when ecological dynamics come into play. Our results, which reflect a generic outcome of biotic interactions in communities, highlight the importance of preserving species that may not seem important for ecosystem functioning based on their effect traits and relative biomass alone.

Our work formalises the fact that, because species interact, their individual contribution to ecosystem functioning is a collective, context-dependent outcome, and that species can dynamically contribute to functions that they do not perform directly. Decades of BEF research have highlighted the context-dependency of species contributions to ecosystem functioning (Sandau et al., 2017; Bannar-Martin et al., 2018), yet using this context dependency to draw conclusions at the species level has rarely been attempted. Sanchez et al. (2023) is a notable exception: using microbial experiments and re-analyzing BEF data, they showed that the effect of introducing a species depends on the level of functioning prior to the introduction. The link we drew between the dynamical contribution and net effects of a species provides a theoretical foundation for the empirical findings of Sanchez et al. (2023).

When a species is a general facilitator or competitor (i.e. the species has a similar effect on other species in the community), the effect of its introduction on ecosystem functioning is incremental: the level of functioning after the introduction of the species cannot be radically different from the functioning prior to its introduction - this dependency has been pointed out in previous work and is analogous to global epistasis and the effect of a mutation (Sanchez et al., 2023; Diaz-Colugna et al., 2024). This finding resonates with those of Liautaud et al. (2019) who show that when interactions exhibit a low variance in the community, a small change in environmental conditions leads to a small change in community composition. On the contrary, when interactions are heterogeneous, the dynamic contribution of species can radically depart from its mean effect. We thus expect the relationship pointed out in Diaz-Colugna et al. (2024) and in Eq. A12 to erode as the analysis is reproduced in organisms that interact with higher variability among each other (eg. a community in which the by-product of one bacteria can be used as a resource for some species while it is toxic to others).

Determining the dynamic contribution of species *in situ* or from experiments is challenging. Estimating the dynamic contribution of species in experimental systems requires particular data: monocultures, effect traits, and ways to estimate interactions (eg. duo-cultures), data from richer communities (eg., data of BEF experiments), or data on monoculture and polyculture sensitivity to environmental perturbations, even if populations have not stabilized (Appendix C). Re-analyzing several BEF experiments with our approach may shed light on the proportions in which different types of species (redundant with a null dynamic contribution, keystone with a strong dynamic contribution) are present in experimental communities. Devising a method that assesses the dynamic contribution of species *in situ* remains an important next step. Long-term temporal series of ecosystem function, response to different types of perturbations, coupled with abundance data are likely needed, in the vein of methods used to estimate biotic interactions (eg. gradient matching, Bonnafé and Coulson, 2023).

Rare species, whose functional role has notoriously been challenging to understand (Lyons et al., 2005), may contribute to ecosystem functions in the dynamic sense. When only considering static contributions, rare species can only matter if they are the only ones to contribute to a narrow function, i.e. they have a unique effect trait. Rare species uniquely contributing to functions have been identified in a variety of ecosystems (eg. Mouillot et al. 2013 in coral reef fishes and tropical trees; Pester et al., 2010 in peatland microbial communities, Soliveres et al., 2016 in grasslands). In these cases, it is obvious that losing a rare species threatens the ecosystem function to which it contributes (O’Gordman et al. 2011). With our approach, we highlight, less intuitively, that rare species can also have a high contribution in the dynamic sense, via their interactions with other species. In plant communities, species of the genus *Equisitum* in wetlands or *Lupinus* in grasslands (Hinsinger et al., 2011; Marsh et al. 2000), for example, could have a high dynamic contribution. These two genera, though not abundant (5% of total biomass for *Equisitum*), increase the availability of phosphorous for *Lupinus* (Hinsinger et al., 2011, Jaillard et al., 2018), phosphorous, potassium and calcium for *Equisitum* (Marsh et al., 2000), thereby facilitating the species that end up being dominant. These rare species, via their interactions, are important in the dynamic sense and qualify as keystone species (Paine et al., 1966). Historically, keystone species have been identified from their direct interactions with other species (predation in the case of Paine et al., 1966). Our results suggest that some rare species that do not directly interact strongly with others may qualify as keystone species but as a collective outcome of the whole community, through via potentially very indirect interactions.

Static and dynamic contributions to ecosystem functioning have different implications for conservation. While conservationists may be tempted, in order to preserve a function or implement a restoration plan, to prioritize species based on their static contribution ecosystem functioning (Laughlin, 2014), the notion of dynamic contribution reveals that focusing on individual traits and biomass can be misleading, and mask the role of some species. Species that interact strongly with others are likely to have strong dynamic contributions (Soulé et al., 2003, Harvey et al., 2017). However, because dynamic contributions are context-dependent, a species that has a high contribution in one system may not necessarily have a high contribution in another. We therefore argue that conservation measures aiming to protect entire communities may be more successful in preserving ecosystem functioning than those targeting single species.

Species’ dynamic contributions are revealed, both formally and in practice, when considering the response of ecosystem functions to press perturbations. Ideally we should consider weak perturbations that modify species’ intrinsic characteristics (eg. a pesticide that reduces growth rates, habitat destruction reducing carrying capacities) without causing secondary extinctions, nor modifying the interactions between species —conditions allowing for a linear theory to apply. In practice, this is a lot to ask: the strength of interactions may vary with the environmental context (eg. temperature change, fertilization; Callaway et al., 2002; Piccardi et al., 2019; Koffel et al., 2018), and secondary extinctions are hard to avoid when considering species-rich systems comprising many rare species, and/or communities behaving like cliques (Liautaud et al., 2019). Formally, species dynamic contributions can still be revealed by perturbations that modify species interactions (see Appendix E). However it is clear that generalizing the two-way relationship between the dynamic contribution and response to perturbations to contexts where the environment undergoes fluctuations, continuous change, or larger shifts in community composition, is an interesting but very challenging next step.

## Conclusion

A major objective of functional ecology is to find measurable traits of organisms that scale-up to the functioning of whole populations (Lavorel and Garnier, 2002; Violle et al., 2007). Our work complements this research agenda by highlighting the role of biotic interactions in shaping the distribution of the biomass across species. In fact, our approach reveals how, by neglecting biotic interactions, the mass ratio-hypothesis and its applications (via community-weighted means) can fail to fully grasp the role played by species in ecosystem functioning. The notion of dynamic contribution of species is particularly relevant in a changing world because, unlike the static contribution, it is designed to reflect the impact of perturbations on ecosystem functions. Our work is merely a proof of concept, but points towards an important and ambitious goal: leverage long-term experiments or observations, monitoring species biomasses after disturbances, to infer the dynamic contribution of species in nature, and reveal their hidden importance for ecosystem functioning.

## References

Arnoldi, Jean-François, Matthieu Barbier, Ruth Kelly, György Barabás, and Andrew L. Jackson. 2022. “Invasions of Ecological Communities: Hints of Impacts in the Invader’s Growth Rate.” Methods in Ecology and Evolution 13 (1): 167–82. 10.1111/2041-210X.13735.

Bannar-Martin, Katherine H., Colin T. Kremer, S.K. Morgan Ernest, Mathew A. Leibold, Harald Auge, Jonathan Chase, Steven A.J. Declerck, et al. 2018. “Integrating Community Assembly and Biodiversity to Better Understand Ecosystem Function: The Community Assembly and the Functioning of Ecosystems (CAFE) Approach.” Ecology Letters 21 (2): 167–80. 10.1111/ele.12895.

Bezanson, Jeff, Alan Edelman, Stefan Karpinski, and Viral B Shah. 2017. “Julia: A Fresh Approach to Numerical Computing.” SIAM Review 59 (1): 65–98.

Brun, Philipp, Cyrille Violle, David Mouillot, Nicolas Mouquet, Brian J. Enquist, François Munoz, Tamara Münkemüller, Annette Ostling, Niklaus E. Zimmermann, and Wilfried Thuiller. 2022. “Plant Community Impact on Productivity: Trait Diversity or Key(Stone) Species Effects?” Ecology Letters 25 (4): 913–25. 10.1111/ele.13968.

Bunin, Guy. 2017. “Ecological Communities with Lotka-Volterra Dynamics.” Physical Review E 95 (4): 042414. 10.1103/PhysRevE.95.042414.

Callaway, Ragan M., R. W. Brooker, Philippe Choler, Zaal Kikvidze, Christopher J. Lortie, Richard Michalet, Leonardo Paolini, et al. 2002. “Positive Interactions among Alpine Plants Increase with Stress.” Nature 417 (6891): 844–48. 10.1038/nature00812.

Cardinale, Bradley J., Justin P. Wright, Marc W. Cadotte, Ian T. Carroll, Andy Hector, Diane S. Srivastava, Michel Loreau, and Jerome J. Weis. 2007. “Impacts of Plant Diversity on Biomass Production Increase through Time Because of Species Complementarity.” Proceedings of the National Academy of Sciences 104 (46): 18123–28. 10.1073/pnas.0709069104.

Delalandre, Léo, Pierre Gaüzère, Wilfried Thuiller, Marc Cadotte, Nicolas Mouquet, David Mouillot, François Munoz, et al. 2022. “Functionally Distinct Tree Species Support Long-Term Productivity in Extreme Environments.” Proceedings of the Royal Society B: Biological Sciences 289 (1967): 20211694. 10.1098/rspb.2021.1694.

Díaz, S. (2025). Plant functional traits and the entangled phenotype. Functional Ecology, 39(5), 1144–1159. 10.1111/1365-2435.70017

Diaz-Colunga, Juan, Abigail Skwara, Jean C.C. Vila, Djordje Bajic, and Alvaro Sanchez. 2024. “Global Epistasis and the Emergence of Function in Microbial Consortia.” Cell 187 (12): 3108-3119.e30. 10.1016/j.cell.2024.04.016.

Enquist, Brian J., Jon Norberg, Stephen P. Bonser, Cyrille Violle, Colleen T. Webb, Amanda Henderson, Lindsey L. Sloat, and Van M. Savage. 2015. “Scaling from Traits to Ecosystems.” In Advances in Ecological Research, 52:249–318. Elsevier. 10.1016/bs.aecr.2015.02.001.

Fant, Lorenzo, and Giulia Ghedini. 2024. “Biomass Competition Connects Individual and Community Scaling Patterns.” Nature Communications 15 (1): 9916. 10.1038/s41467-024-54307-w.

Fargione, Joseph, David Tilman, Ray Dybzinski, Janneke Hille Ris Lambers, Chris Clark, W. Stanley Harpole, Johannes M.H Knops, Peter B Reich, and Michel Loreau. 2007. “From Selection to Complementarity: Shifts in the Causes of Biodiversity–Productivity Relationships in a Long-Term Biodiversity Experiment.” Proceedings of the Royal Society B: Biological Sciences 274 (1611): 871–76. 10.1098/rspb.2006.0351.

Garnier, Eric, Jacques Cortez, Georges Billès, Marie-Laure Navas, Catherine Roumet, Max Debussche, Gérard Laurent, et al. 2004. “Plant Functional Markers Capture Ecosystem Properties During Secondary Succession.” Ecology 85 (9): 2630–37. 10.1890/03-0799.

Grime, J. P. 1998. “Benefits of Plant Diversity to Ecosystems: Immediate, Filter and Founder Effects.” Journal of Ecology 86 (6): 902–10. 10.1046/j.1365-2745.1998.00306.x.

Harvey, Eric, Isabelle Gounand, Colette L. Ward, and Florian Altermatt. 2017. “Bridging Ecology and Conservation: From Ecological Networks to Ecosystem Function.” Journal of Applied Ecology 54 (2): 371–79. 10.1111/1365-2664.12769.

Heilpern, Sebastian A., Brian C. Weeks, and Shahid Naeem. 2018. “Predicting Ecosystem Vulnerability to Biodiversity Loss from Community Composition.” Ecology 99 (5): 1099–1107. 10.1002/ecy.2219.

Hinsinger, Philippe, Elodie Betencourt, Laetitia Bernard, Alain Brauman, Claude Plassard, Jianbo Shen, Xiaoyan Tang, and Fusuo Zhang. 2011. “P for Two, Sharing a Scarce Resource: Soil Phosphorus Acquisition in the Rhizosphere of Intercropped Species.” Plant Physiology 156 (3): 1078–86. 10.1104/pp.111.175331.

Hong, Pubin, Bernhard Schmid, Frederik De Laender, Nico Eisenhauer, Xingwen Zhang, Haozhen Chen, Dylan Craven, et al. 2022. “Biodiversity Promotes Ecosystem Functioning despite Environmental Change.” Ecology Letters 25 (2): 555–69. 10.1111/ele.13936.

Hooper, D. U., F. S. Chapin, J. J. Ewel, A. Hector, P. Inchausti, S. Lavorel, J. H. Lawton, et al. 2005. “Effects of Biodiversity on Ecosystem Functioning: A Consensus of Current Knowledge.” Ecological Monographs 75 (1): 3–35. 10.1890/04-0922.

Intergovernmental Science-Policy Platform on Biodiversity and Ecosystem Services, IPBES. 2019. Summary for Policymakers of the Global Assessment Report on Biodiversity and Ecosystem Services. IPBES secretariat, Bonn, Germany. 10.5281/ZENODO.3553579.

Jaillard, Benoît, Camille Richon, Philippe Deleporte, Michel Loreau, and Cyrille Violle. 2018. “An a Posteriori Species Clustering for Quantifying the Effects of Species Interactions on Ecosystem Functioning.” Methods in Ecology and Evolution 9 (3): 704–15. 10.1111/2041-210X.12920.

Koffel, Thomas, Simon Boudsocq, Nicolas Loeuille, and Tanguy Daufresne. 2018. “Facilitation-vs. Competition-Driven Succession: The Key Role of Resource-Ratio.” Ecology Letters 21 (7): 1010– 21. 10.1111/ele.12966.

Lajaaiti, Ismaël, Sonia Kéfi, Michel Loreau, Alice Ardichvili, and Jean-François Arnoldi. 2025. “Revealing the Organization of Species Stability in Ecological Communities.” BioRXiv 10.1101/2025.03.25.645234.

Laughlin, Daniel C. 2014. “Applying Trait-based Models to Achieve Functional Targets for Theory-driven Ecological Restoration.” Ecology Letters 17 (7): 771–84. 10.1111/ele.12288.

Lavorel, S., and E. Garnier. 2002. “Predicting Changes in Community Composition and Ecosystem Functioning from Plant Traits: Revisiting the Holy Grail: Plant Response and Effect Groups.” Functional Ecology 16 (5): 545–56. 10.1046/j.1365-2435.2002.00664.x.

Law, Richard, and R. Daniel Morton. 1996. “Permanence and the Assembly of Ecological Communities.” Ecology 77 (3): 762–75. 10.2307/2265500.

Leitão Rafael P., Jansen Zuanon, Sébastien Villéger, Stephen E. Williams, Christopher Baraloto, Claire Fortunel, Fernando P. Mendonça, and David Mouillot. 2016. “Rare Species Contribute Disproportionately to the Functional Structure of Species Assemblages.” Proceedings of the Royal Society B: Biological Sciences 283 (1828): 20160084. 10.1098/rspb.2016.0084.

Liautaud, Kevin, Egbert H. Van Nes, Matthieu Barbier, Marten Scheffer, and Michel Loreau. 2019. “Superorganisms or Loose Collections of Species? A Unifying Theory of Community Patterns along Environmental Gradients.” Ecology Letters 22 (8): 1243–52. 10.1111/ele.13289.

Loreau, Michel, and Andy Hector. 2001. “Partitioning Selection and Complementarity in Biodiversity Experiments.” Nature 412 (6842): 72–76. 10.1038/35083573.

Loreau, M., Naeem, S., Inchausti, P., Bengtsson, J., Grime, J. P., Hector, A., Hooper, D. U., Huston, M. A., Raffaelli, D., Schmid, B., Tilman, D., & Wardle, D. A. 2001. “Biodiversity and Ecosystem Functioning: Current Knowledge and Future Challenges.” Science 294(5543): 804– 808. 10.1126/science.1064088.

Loreau, Michel, Andy Hector, and Forest Isbell. 2022. The Ecological and Societal Consequences of Biodiversity Loss. Sciences. Ecosystems and Enviroment. Biodiversity. London: ISTE Ltd.

Lyons, K. G., C. A. Brigham, B.H. Traut, and M. W. Schwartz. 2005. “Rare Species and Ecosystem Functioning.” Conservation Biology 19 (4): 1019 – 24. 10.1111/j.1523-1739.2005.00106.x.

Marsh, Anne S., John A. Arnone, Bernard T. Bormann, and John C. Gordon. 2000. “The Role of Equisetum in Nutrient Cycling in an Alaskan Shrub Wetland.” Journal of Ecology 88 (6): 999–1011. 10.1046/j.1365-2745.2000.00520.x.

Mouillot, David, David R. Bellwood, Christopher Baraloto, Jerome Chave, Rene Galzin, Mireille Harmelin-Vivien, Michel Kulbicki, et al. 2013. “Rare Species Support Vulnerable Functions in High-Diversity Ecosystems.” PLoS Biology 11 (5): e1001569. 10.1371/journal.pbio.1001569.

O’Gorman Eoin J., Jon M. Yearsley, Tasman P. Crowe, Mark C. Emmerson, Ute Jacob, and Owen L. Petchey. 2011. “Loss of Functionally Unique Species May Gradually Undermine Ecosystems.” Proceedings of the Royal Society B: Biological Sciences 278 (1713): 1886–93. 10.1098/rspb.2010.2036.

Paine, Rt. 1966. “Food Web Complexity and Species Diversity.” American Naturalist 100 (910): 65-+. 10.1086/282400.

Peltzer, Duane A, Peter J Bellingham, Hiroko Kurokawa, Lawrence R Walker, David A Wardle, and Gregor W Yeates. 2009. “Punching above Their Weight: Low-Biomass Non-Native Plant Species Alter Soil Properties during Primary Succession.” Oikos 118:1001–14. 10.1111/j.1600-0706.2009.17244.x.

Pennekamp, Frank, Mikael Pontarp, Andrea Tabi, Florian Altermatt, Roman Alther, Yves Choffat, Emanuel A. Fronhofer, et al. 2018a. “Biodiversity Increases and Decreases Ecosystem Stability.” Nature 563 (7729): 109–12. 10.1038/s41586-018-0627-8.

Pennekamp, Frank, Mikael Pontarp, Andrea Tabi, Florian Altermatt, Roman Alther, Yves Choffat, Emanuel Fronhofer, et al. 2018b. “Pennekampster/Code_and_data_OverallEcosystemStability: Release of Data and Code.” Zenodo. 10.5281/ZENODO.1345556.

Pester, Michael, Norbert Bittner, Pinsurang Deevong, Michael Wagner, and Alexander Loy. 2010. “A ‘Rare Biosphere’ Microorganism Contributes to Sulfate Reduction in a Peatland.” The ISME Journal 4 (12): 1591–1602. 10.1038/ismej.2010.75.

Piccardi, Philippe, Björn Vessman, and Sara Mitri. 2019. “Toxicity Drives Facilitation between 4 Bacterial Species.” Proceedings of the National Academy of Sciences 116 (32): 15979–84. 10.1073/pnas.1906172116.

Pichon, Benoît, Sonia Kéfi, Nicolas Loeuille, Ismaël Lajaaiti, and Isabelle Gounand. 2024. “Integrating Ecological Feedbacks across Scales and Levels of Organization.” Ecography, April, e07167. 10.1111/ecog.07167.

Rackauckas, Christopher, and Qing Nie. 2017. “DifferentialEquations.Jl–a Performant and Feature-Rich Ecosystem for Solving Differential Equations in Julia.” Journal of Open Research Software 5 (1).

Sanchez, Alvaro, Djordje Bajic, Juan Diaz-Colunga, Abigail Skwara, Jean C.C. Vila, and Seppe Kuehn. 2023. “The Community-Function Landscape of Microbial Consortia.” Cell Systems 14 (2): 122–34. 10.1016/j.cels.2022.12.011.

Sanchez-Gorostiaga, Alicia, Djordje Bajić, Melisa L. Osborne, Juan F. Poyatos, and Alvaro Sanchez. 2019. “High-Order Interactions Distort the Functional Landscape of Microbial Consortia.” PLOS Biology 17 (12): e3000550. 10.1371/journal.pbio.3000550.

Sandau, Nadine, Yvonne Fabian, Odile T. Bruggisser, Rudolf P. Rohr, Russell E. Naisbit, Patrik Kehrli, Alexandre Aebi, and Louis-Félix Bersier. 2017. “The Relative Contributions of Species Richness and Species Composition to Ecosystem Functioning.” Oikos 126 (6): 782–91. 10.1111/oik.03901.

Smith, Melinda D., Sally E. Koerner, Alan K. Knapp, Meghan L. Avolio, Francis A. Chaves, Elsie M. Denton, John Dietrich, et al. 2020. “Mass Ratio Effects Underlie Ecosystem Responses to Environmental Change.” Journal of Ecology 108 (3) : 855–64. 10.1111/1365-2745.13330.

Soliveres, Santiago, Peter Manning, Daniel Prati, Martin M. Gossner, Fabian Alt, Hartmut Arndt, Vanessa Baumgartner, et al. 2016. “Locally Rare Species Influence Grassland Ecosystem Multifunctionality.” Philosophical Transactions of the Royal Society B: Biological Sciences 371 (1694): 20150269. 10.1098/rstb.2015.0269.

Soulé Michael E., James A. Estes, Joel Berger, and Carlos Martinez Del Rio. 2003. “Ecological Effectiveness: Conservation Goals for Interactive Species.” Conservation Biology 17 (5): 1238–50. 10.1046/j.1523-1739.2003.01599.x.

Violle, Cyrille, Marie-Laure Navas, Denis Vile, Elena Kazakou, Claire Fortunel, Irène Hummel, and Eric Garnier. 2007. “Let the Concept of Trait Be Functional!” Oikos 116 (5): 882–92. 10.1111/j.0030-1299.2007.15559.x.

Wasof, Safaa, Jonathan Lenoir, Tarek Hattab, Aurélien Jamoneau, Emilie Gallet-Moron, Evy Ampoorter, Robert Saguez, et al. 2018. “Dominance of Individual Plant Species Is More Important than Diversity in Explaining Plant Biomass in the Forest Understorey.” Journal of Vegetation Science 29 (3): 521–31. 10.1111/jvs.12624.

Zelnik, Yuval R., Nuria Galiana, Matthieu Barbier, Michel Loreau, Eric Galbraith, and Jean-François Arnoldi. 2024. “How Collectively Integrated Are Ecological Communities?” Ecology Letters 27 (1): e14358. 10.1111/ele.14358.

