## Supplementary Material A-E for "Beyond biomass: how interactions shape species’ role for ecosystem functioning"

### Appendix A: from net effects to dynamic contributions, and relationship with background functioning

We start from our definition of the net effect of species *i* on ecosystem functioning:

$$\Delta\Phi_i = \Phi - \Phi^{\backslash i} \qquad \text{A1}$$

Noting $\delta N_j = N_j - N_{j/i}$ the change in the biomass of species *j* as a consequence of the presence of species *i*, where $\delta N_i = N_i$ by definition, the net effect on functioning reads:

$$\Delta\Phi_i = \sum_{j=1}^{S} t_{Ej}\delta N_j \qquad \text{A2}$$

Removing species *i* amounts to perturbing its carrying capacity enough to push the species to extinction. Assuming linearity, this perturbation $-\delta K_i$ can be computed as

$$\delta N_i = \frac{\partial N_i}{\partial K_i}\delta K_i = V_{ii}\delta K_i = N_i \qquad \text{A3}$$

This specific perturbation impacts other species as well, and at linear order:

$$\delta N_j = V_{ji}\delta K_i = V_{ji}K_i^* \times \frac{N_i/K_i^*}{V_{ii}} \qquad \text{A4}$$

We can then obtain the change in ecosystem functioning resulting from changes in the biomass of all species where we recognize the expression of $\Phi_i^{dyn}$ from its derivation Eq. 4 of the main text.:

$$\Delta\Phi_i = \frac{N_i/K_i^*}{V_{ii}}\sum_{j=1}^{S} t_{Ej}V_{ij}K_i^* = \frac{N_i/K_i^*}{V_{ii}}\Phi_i^{dyn} \qquad \text{A5}$$

We can manipulate the expression of $\Phi_i^{dyn}$ further to obtain another expression in which the background level of functioning $\Phi^{\backslash i}$ appears. The idea is to note that species *i* acts as a perturbation on the community formed without it. This perturbation directly amounts to a change $\delta K_k = A_{ki}N_i$ of the carrying capacity of any species *k*. Thus, noting $a_{ki} = A_{ki}/K_{k/i}^*$ , we may write, still assuming linearity, that the shift in biomass of any given species *j* is:

$$\delta N_j = \sum_{k \neq i} \frac{\partial N_{j/i}}{\partial K_k} K_{k/i}^* a_{ki} N_i \qquad \text{A6}$$

With that in mind, we can now note that

$$\Phi_i^{dyn} = t_{E_i}\frac{\partial N_i}{\partial K_i}K_i^* + \sum_{j \neq i} t_{E_j}\frac{\partial N_{j/i}}{\partial K_i}K_i^* + \sum_{j \neq i} t_{E_j}\frac{\partial \delta N_j}{\partial K_i}K_i^* \qquad \text{A7}$$

The second term in the r.h.s. vanishes because $N_{j/i}$ does not depend on $K_i$. For the last term we can use the expression of $\delta N_j$ just derived. We get to:

$$\sum_{j\neq i} t_{E_j}\frac{\partial \delta N_j}{\partial K_i}K_i^* = \frac{\partial N_i}{\partial K_i}K_i^* \sum_{k,j\neq i} t_{E_j}\frac{\partial N_{j/i}}{\partial K_k}K_{k/i}^* a_{ki} = \frac{\partial N_i}{\partial K_i}K_i^* \sum_{k\neq i} \Phi_{k/i}^{dyn} a_{ki} \quad \text{A8}$$

Where we recognized the dynamical contributions of species in the community *without* species *i.* If we denote $\Phi_i^* = t_{E_i}K_i^*$, which we may think of as the functioning of a monoculture of *i*, then we get to

$$\Phi_i^{dyn} = V_{ii}(\Phi_i^* + \sum_{k\neq i} \Phi_{k/i}^{dyn} a_{ki} K_i^*) \quad \text{A9}$$

Because $\sum_{k\neq i} \Phi_{k/i}^{dyn} = \Phi_{/i}$ —the background functioning of the community without species *i*— then we may simplify the expression further and recognize Eq. 8 in the main text:

$$\Phi_i^{dyn} = V_{ii}(\Phi_i^* + \Phi_{/i}\overline{a}_i K_i^*) \quad \text{A10}$$

Where the term $\overline{a}_i K_i^*$ can be interpreted as the weighted mean of the interactions incoming from species *i*. This weighted mean has to be understood with care since the weights are the relative dynamical contributions of affected species, and those contributions could be negative. This will not matter if the interactions $a_{ki}$ are uniform, but could possibly reverse some expectations, making an apparently facilitative species have a negative effect on functioning if it facilitates a strongly antagonistic species (i.e. whose dynamical contribution is negative).

We can go back to the expression of the net effect $\Phi_i^{dyn}\eta_i / V_{ii}$ and deduce:

$$\Delta\Phi_i = t_{E_i}N_i + \sum_{k\neq i} \Phi_{k/i}^{dyn} a_{ki} N_i \quad \text{A11}$$

Because $\sum_{k\neq i} \Phi_{k/i}^{dyn} = \Phi_{/i}$, we can then simplify:

$$\Delta\Phi_i = \Phi_i^{stat} + \Phi_{/i}\overline{a}_i N_i \quad \text{A12}$$

The dependence between the response of ecosystem functioning and the background functioning has been empirically found in microbial communities (Sanchez et al., 2023).

### Appendix B: estimating dynamic contributions from net effects

Using simulations, we assessed whether the dynamic contribution of species could be estimated from Eq. 11, making the simplifying assumptions that the pseudo relative yield $\eta_i^* = N_i/K_i^*$ is close to the actual relative yield $\eta_i = N_i/K_i$, and that the net auto-interaction effect is uniform:

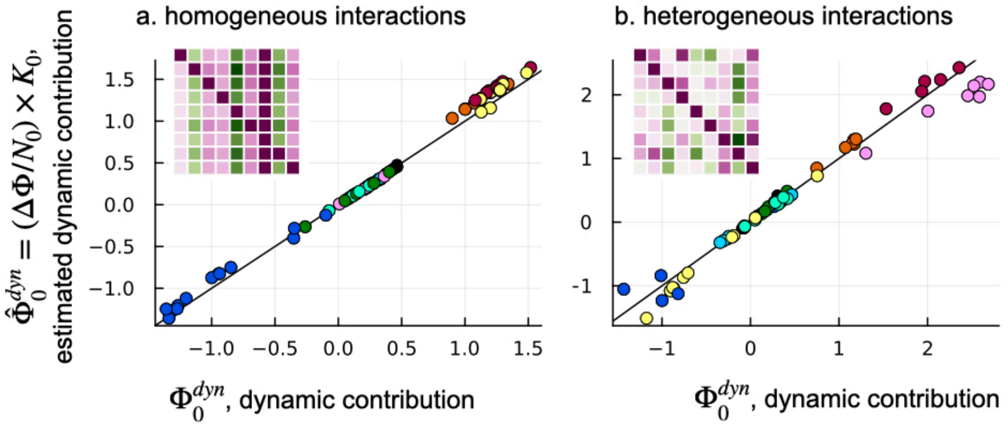


**Figure B1: the dynamic contributions of a species $\Phi_0^{dyn} = \sum_j t_{Ej} V_{j0} K_0^*$ (exact expression from Eq. 5) can be estimated from the effect of introducing it in the community $\Delta\Phi_0$ and its relative yield $\eta_0$ . The prediction is accurate whether species have an homogeneous effect on others ($u_a$ = 0) or an heterogeneous effect ($u_a$ = 0.6).**

Simulations show that the change in ecosystem functioning and the relative yield can be used to estimate the dynamic contribution of species, using Eq. 11 (Fig. B1). The relationship holds when the effect of species on others is homogeneous or heterogeneous: see how, in the interaction matrix in the top left of the panels, the columns are of the same color indicating a uniform effect of the column species on others in a, while in b, the same species can have both positive and negative effects on others, and of strongly varying magnitudes.

### Appendix C: estimating the dynamic contribution in out-of-equilibrium dynamics

By defining $K_i^*$ such that $\sum_{i=1}^{S} V_{ji} K_i^* = N_j$, we are not restricted to equilibrium conditions. Wether $K_i^*$ corresponds to the actual carrying capacity $K_i$ of the species (its equilibrium abundance in monoculture) tells us whether the system is at equilibrium. Fig 1D shows that even in out-of equilibrium conditions, the dynamic contribution of species accurately reflects how an ecosystem function changes with the extinction of a species.

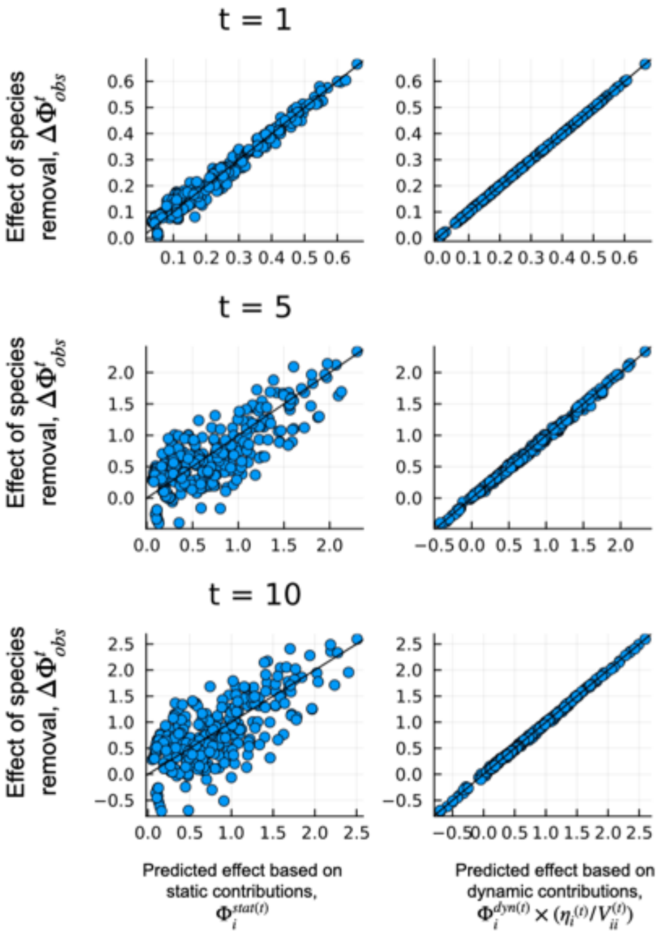


**Fig.C1: In these figures, each point is a comparison of two out of equilibrium communities, that serves as a proxy to estimate the effect of removing one species. The dynamic contribution of species - estimated from out-of-equilibrium duocultures and monocultures- is always better at predicting the effect of removing the species than the static contribution. These plots may seem contradictory with Appendix C: in fact, in Appendix C, only one of the community (the one in which the species was removed) is out of equilibrium. Here, the two communities that are compared to assess the effect of removing a species (as in the BEF data) are out of equilibrium.**

### Appendix D: The relationship between dynamic contributions and effect of removing a species erodes with secondary extinctions

In the main text, Eq. 8 means that the dynamic contribution of species encodes the linear response of the function to a small perturbation of magnitude $\delta K_i$. When the perturbation is the extinction of a species, $\delta K_i = -N_i/V_{ii}$, the perturbation is not small but we assume that the relationship between a species' abundance and the carrying capacities of other species is still linear: $\sum_{i=1}^{S} V_{ji}K_i^* = N_j$. In the simulations of a GLV, this assumption always holds as long as the removal of a species does not cause secondary extinctions, meaning that the matrix of net effects $V$ remains constant. In theory, this means that the relationship between the dynamic contribution of a species and the effect of removing it (Eq. 8) should erode when secondary extinctions occur. However, fig. D1 shows that the dynamic contribution of species is still a good predictor of the effect of its removal, even when it generates secondary extinctions. This is probably because species that go extinct secondarily have a low abundance $N_i$ in the full community.

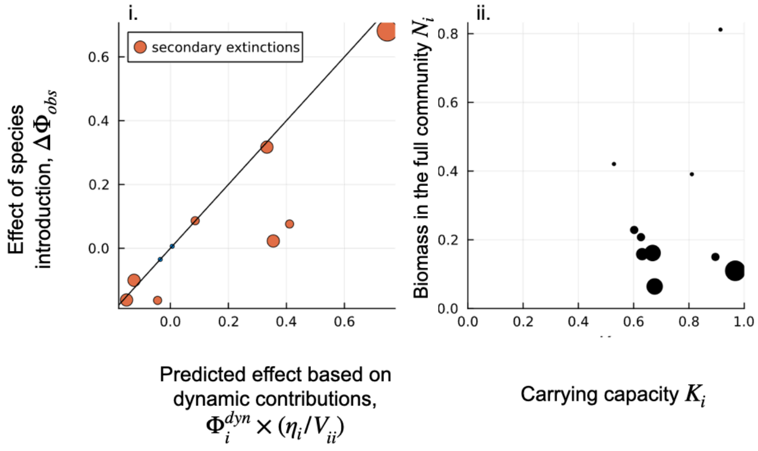


**Figure D1: i shows the predicted effect of introducing a species based on its dynamic contribution. The blue dots correspond to situations where no secondary extinctions occur (the dots fall perfectly on the 1:1 and the orange dots to the situations where secondary extinctions occur. The size of the point increases with the number of secondary extinctions caused. While secondary extinctions imply that the dynamic contribution of species does not perfectly predict the effect of removing it, it still provides a good prediction. In ii., each point corresponds to a species plotted on a plane capturing its carrying capacity and abundance in the community. The size of the point grows with the frequency at which the species got secondarily extinct.**

### Appendix E: The dynamic contribution of species with varying interactions

In this appendix, we expand our framework to a case where environmental changes affects both the intrinsic characteristics of species and the strength of interactions. For simplicity we focus on Lotka-Volterra models at equilibrium but the reasoning is more general:

$$N(E) = V(E)K^*(E)$$

$$\frac{\partial N}{\partial E} = \frac{\partial V}{\partial E}K^* + V\frac{\partial K^*}{\partial E}$$

$$\frac{\partial V}{\partial E} = V\frac{\partial A}{\partial E}V \text{ where } -A = V^{-1}$$

$A(E)$ is the pairwise matrix of direct interactions that depends on environmental conditions $E$

$$\frac{\partial N}{\partial E} = V\frac{\partial A}{\partial E}VK^* + V\frac{\partial K^*}{\partial E}$$

$$\frac{\partial N}{\partial E} = V\left(\frac{\partial A}{\partial E}N + \frac{\partial K^*}{\partial E}\right) \text{ because } VK^* = N$$

$$\frac{\partial N_i}{\partial E} = \sum_{j=1}^{S} V_{ij}K_j^*\left(\frac{\sum_{k=1}^{S}\frac{\partial A_{jk}}{\partial E}N_k + \frac{\partial K_j^*}{\partial E}}{K_j^*}\right)$$

The resulting change in ecosystem functioning reads:

$$\sum_{i=1}^{S} t_{E_i}\frac{\partial N_i}{\partial E} = \sum_{i,j=1}^{S} t_{E_i}V_{ij}K_j^*\left(\frac{\sum_{k=1}^{S}\frac{\partial A_{jk}}{\partial E}N_k + \frac{\partial K_j^*}{\partial E}}{K_j^*}\right)$$

We see that the dynamic contribution appears:

$$\frac{d\Phi}{dE} = \sum_{j}\Phi_j^{dyn}\left(\frac{\sum_k\frac{\partial A_{jk}}{\partial E}N_k + \frac{\partial K_j^*}{\partial E}}{K_j^*}\right) \qquad \text{(E1)}$$

Equation E1 states that the dynamic contributions of species drive the response of an ecosystem function to environmental change even in the case of varying interactions. The dynamic contribution needs to be weighted by the intrinsic sensitivity of the species to the environmental change $\left(\frac{\partial K_i}{\partial E}/K_i^*\right)$, but also by how the direct effects of other species have changed with environmental change. The equality in E1 is illustrated in Fig. E1c.

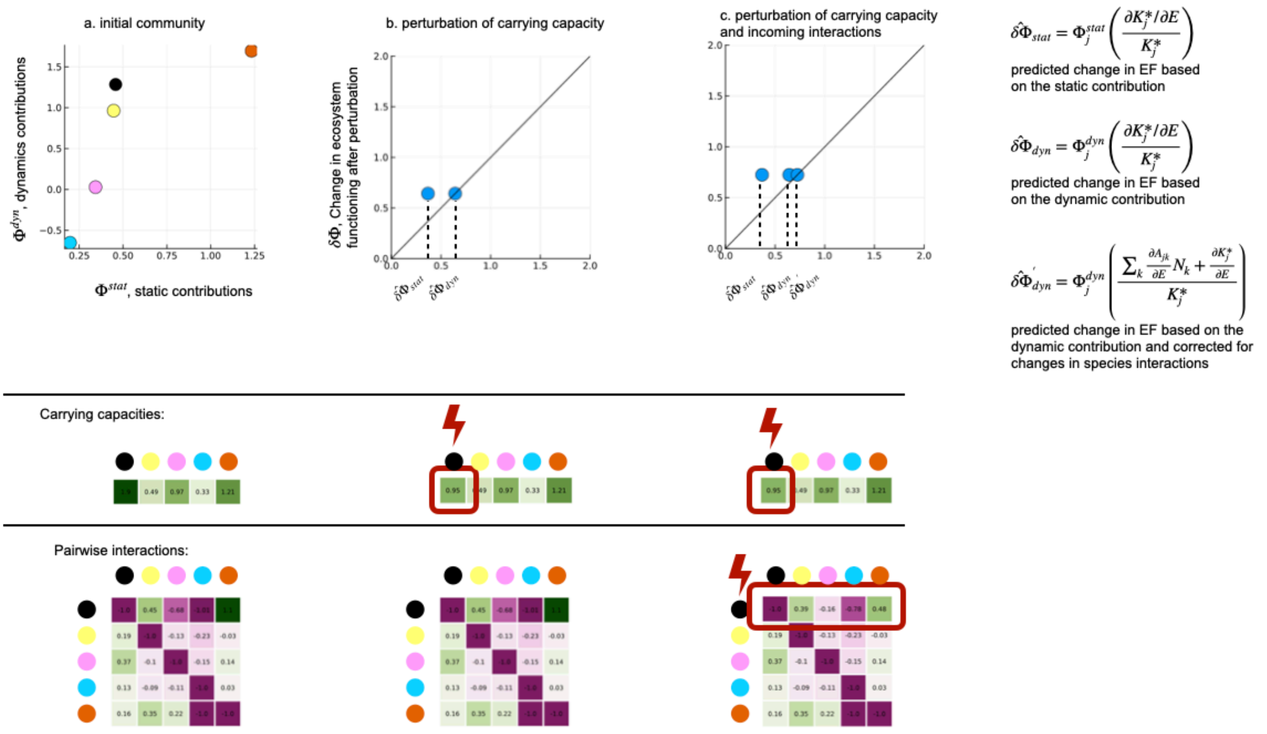


**Figure E1: Simulated example of a perturbation that affects both carrying capacities and pairwise interactions. a. Community of 5 species with randomly drawn interaction terms and carrying capacities. The black species is relatively rare with a strong dynamic contribution. b. The carrying capacity of the black species is decreased by 50% and the response of the ecosystem function (EF) reflects the dynamic contribution of the species. c. The carrying capacity of the black species is decreased by 50% and incoming interactions on the black species are decreased at random (perturbation randomly drawn between 0 and 1).**
